# Extracellular methylglyoxal; the passage across brain endothelial cells and the effect on barrier function

**DOI:** 10.64898/2026.02.20.706964

**Authors:** Eline Berends, Sarah Guimbal, Xiaodi Zhang, Isa Frijns, Marjo P.H. van de Waarenburg, Jean L.J.M. Scheijen, Kristiaan Wouters, Robert J. van Oostenbrugge, Britta Engelhardt, Casper G. Schalkwijk, Sébastien Foulquier

**Author notes:** **Corresponding author:** Prof Dr Casper Schalkwijk, Maastricht University, Faculty of Health Medicine and Life Sciences, Department of Internal Medicine, Universiteitssingel 50, 6229 ER Maastricht, The Netherlands. Shared last-authorship (Casper G. Schalkwijk, Sébastien Foulquier).

## Abstract

**Background:** Methylglyoxal (MGO), a highly reactive by-product of glycolysis, has been associated with cognitive decline and Alzheimer’s disease, though the mechanistic role of MGO remains unclear. Moreover, conflicting findings exist regarding MGO’s toxicity on the blood-brain barrier (BBB). This study investigated whether MGO can cross the BBB under physiologically relevant conditions and whether MGO affects BBB permeability.

**Methods:** Mice were intravenously injected with highly purified home-made MGO or PBS, and MGO concentration was measured at five timepoints in the cerebral cortex up to 4 hours after injection. MGO toxicity was screened on a human brain endothelial cell line (hCMEC/D3) using a live/dead assay prior to the study of selected MGO concentrations and on hiPSC-derived brain microvascular endothelial cells (EECM-BMECs). EECM-BMECs were cultured on Transwell® inserts, and barrier function was assessed by sodium fluorescein permeability and transendothelial passage of ^13^C3-MGO quantified by UPLC-MS/MS.

**Results:** MGO levels in the mouse cortex did not increase post-injection. MGO was not toxic to hCMEC/D3 cells, and it had no impact on barrier properties of EECM-BMECs. After 1-hour exposure, ∼13% of total ^13^C3-MGO was recovered in its free form, and only ∼1% of supplemented MGO was recovered from the abluminal side.

**Conclusion:** MGO does not cross the BBB in vivo and does not affect barrier properties of a human in vitro model of the BBB. In vitro MGO passage across the BBB is minimal. These findings suggest that circulating MGO is unlikely to directly affect neuronal function via BBB disruption or enter the brain in its free from.

## Introduction

Methylglyoxal (MGO) is a highly reactive by-product of glycolysis and a major precursor in the formation of advanced glycation endproducts. It has been implicated in many diseases including diabetes and its vascular complications [1] as well as brain related disorders [2–4]. Several studies reported an association between elevated plasma levels of MGO with cognitive impairment in elderly individuals and people with Alzheimer’s disease [5–7]. It remains, however, unclear whether MGO from the circulation can damage and/or directly cross the blood-brain barrier (BBB) to exert detrimental effects on the brain.

The brain vasculature is highly specialised, by presence of the BBB, which is essential in protecting the brain from toxins and pathogens and maintaining a tightly regulated environment for neuronal function [8]. The BBB consists of endothelial cells, astrocytes, and mural cells, including pericytes and vascular smooth muscle cells [8, 9]. Preserving the BBB is essential for maintaining healthy neuronal functioning in the brain, and damage to one or more of its components will have major consequences for brain function. Vascular disfunction has been implicated in several neurodegenerative disorders [10].

Various studies have reported that MGO is toxic to brain microvascular endothelial cells, and it has been shown that MGO increases barrier permeability and might form a threat for BBB integrity by damaging endothelial cells and pericytes [11]. However, many of these findings are based on studies using commercially available MGO, which is known to be contaminated with formaldehydes and other contaminants that may confound results [12]. Moreover, our recent in vivo work demonstrated that increasing levels of MGO in plasma in young healthy mice, to concentrations comparable to those seen in people with type 2 diabetes, has no consequence for cognitive function, BBB integrity, nor neurovascular coupling response [13].

These apparently conflicting findings raised the question whether circulating MGO can affect BBB properties or even cross the BBB and thereby directly affect neuronal functioning. Therefore, the aim of this study was to investigate whether MGO affects BBB properties and/or crosses the BBB. To study this, we made use of highly purified home-made MGO throughout the study.

## Materials and methods

### Methylglyoxal synthesis

MGO was synthesized in-house by hydrolyzing methylglyoxal-1,1-dimethyl acetal with sulfuric acid, followed by purification through fractional distillation under reduced pressure in a nitrogen atmosphere [14, 15].

### Animal experiments

All animal work was performed in accordance with the EU directive 2010/63/EU, the ARRIVE guidelines and approved by the Dutch Central Committee Animal Experiments and the Animal Welfare Body of the University of Maastricht under permit AVD1070020187086. Male C57Bl/6J mice (Charles River) of 11-12 weeks of age were used for these experiments. Animals were socially housed at temperature and humidity-controlled conditions with a 12-hour light/dark cycle and had ad libitum access to food and water.

Mice received 25 µg of MGO in 200uL saline (MGO) or 200 µL saline (control) injection via the tail vein. Mice were euthanised by CO_2_/O_2_ mixture at 10 minutes (control n=5, MGO n=8), 30 minutes (control n=3, MGO n=5), 1 hour (control n=3, MGO n=5), or 4 hours (control n=3, MGO n=5). The MGO dose was chosen based on a previous study [16]. After sacrifice the cerebral cortex was isolated and snap frozen. The brain tissue was homogenised with a pestle and mortar submerged in liquid nitrogen. The brain homogenate was dissolved in digestion buffer (0.1 M sodium phosphate buffer with 0.02% Triton-X and protease inhibitor [Roche]). After one cycle of freezing and thawing, samples were centrifuged, and protein-containing supernatant was collected for UPLC-MS/MS analysis.

### hCMEC/D3 cells

Human cerebral microvascular endothelial cell line D3 (hCMEC/D3, passage 39 to 41) was maintained at 37°C, 5% CO_2_, in Cultrex® (R&D systems, #3443-100-01) coated flasks in endothelial cell basal medium (EBM™-2) (Lonza, #190860) containing 5% heat inactivated fetal bovine serum (iFBS) (TICO, S0115), 1% penicillin-streptomycin (Thermo Fischer Scientific, #15070), 1.4 µM hydrocortisone (Sigma-Aldrich, #H0135), 5 μg/ml ascorbic acid (Sigma-Aldrich, #A4544), 1% chemically defined lipid concentrate (Thermo Fischer Scientific #11905031), 10 mM HEPES (Thermo Fischer Scientific, #15630), and 1 ng/mL fibroblast growth factor 2 (bFGF) (Sigma-Aldrich, #F0291). Medium was changed every 2 days. Upon 90-100% confluence, cells were passaged through trypsinisation using 0.25% (w/v) Trypsin – 0.53mM EDTA solution in PBS.

Cytotoxicity of MGO on hCMEC/D3 cells was determined with a live/dead cell viability imaging kit (ReadyProbes™ Cell Viability Imaging Kit, Thermo Fischer Scientific, #R37609). Cells were seeded in a Cultrex® coated 96-well plate at 40×10^3^ cells/well. After 24 hours, medium was replaced with EBM™-2 medium without iFBS. After 4 hours, cells were incubated with NucGreen® and NucBlue® for 15 minutes at room temperature following manufacturer’s instructions. The cells were imaged with the ImageXpress® Pico Automated Cell Imaging System (Molecular Devices, LLC, San Jose, CA, USA) before MGO exposure (time point 0). Cells were then treated with MGO ranging from 100 nM to 10 mM (90% EBM-2 serum free growth medium, 10% H_2_O containing MGO), control (H_2_O), or positive control (10% DMSO). Cells were imaged again at 30 minutes, and at 1, 2, 4 and 6 hours of exposure. The relative number of dead cells (%) was calculated as the number of dead cells (positive for NucGreen®) divided by the total number of nuclei (positive for NucBlue®).

### hiPSC derived EECM-BMECs

Brain microvascular endothelial (BMEC)-like cells were acquired using the extended endothelial cell culture method (EECM) for the differentiation of human induces pluripotent stem cells (hiPSC) using iPS(IMR90)-4 (WiCell, RRID:CVCL_C437) exactly as described previously [17]. At passage 3 to 5, EECM-BMECs were seeded into collagen/fibronectin coated 12-well Transwell filter (0.4 μm, 12 mm, Corning, #3401) at 112×10^3^ cells per insert in hECSR medium. Medium was changed every 2 days.

Permeability assay: on day 6, medium from the top chamber was discarded and inserts were placed in a new 12-well plate containing 1.5 mL warm hECSR medium per well. 500 μL of hECSR medium containing 1:1000 sodium fluorescein (NaFl) and 2% H_2_O containing 1 or 10 µM MGO or not, was added to the top chamber. The plate was placed in the incubator (37°C, 5% CO_2_). For the consecutive 2 hours, medium from the bottom compartment was collected every 15 minutes. Fluorescent intensity was measured using a fluorescent plate reader (485 nm excitation / 530 nm emission). The clearance volume and permeability (Pe) were calculated as described previously [17]. Briefly, the clearance volume was calculated as the fluorescent signal multiplied by the volume of the bottom chamber, divided by the fluorescent signal in the top chamber at 120 minutes. The permeability is it the linear slope of the clearance volume over time, minus the clearance volume of a blank Transwell insert.

Labelled MGO exposure: on day 6, medium was replaced 4 hours prior to MGO exposure. For the top compartment 490 μL of hECSR medium was used, for the bottom compartment 1.5 mL. After 4 hours, 10 μL of MGO solution in H_2_O (2% of total volume) was added to the top compartment so that the final concentration in the top compartment was either 1 or 10 μM carbon isotope labelled MGO (^13^C_3_-MGO, Toronto Research Chemicals, TRC-P998872). After 1 hour incubation at 37°C, medium from the top and bottom compartment was collected and stored at -20°C until UPLC-MS/MS measurement. The cell inserts were washed once with PBS, then 150 μL NaP buffer with 0.02% Triton-X and protease inhibitor was added and the inserts were frozen overnight. The next day, cells were scraped form the insert and lysates were collected and stored at -20°C until UPLC-MS/MS measurement.

### UPLC-MS/MS

MGO was measured in mouse cortex, cell lysate, and culture medium using UPLC-MS/MS as described previously [13, 18]. ^13^C_3_-MGO data were corrected for a difference in mass spectrometry response factor between ^13^C_3_-MGO and unlabelled MGO (^12^C_3_-MGO), as described previously [19].

### Statistics and data presentation

All statistics were performed using GraphPad Prism (version 10.4.1, GraphPad Software, San Diego). For data measured over time, a two-way ANOVA was used followed by a Tukey’s or Dunnett’s multiple comparison, for other data a one-way ANOVA was used followed by a Tukey’s multiple comparison test. All data are shown as mean ± standard deviation (SD) calculated from the experimental replicates. Data from EECM-BMECs are shown as mean ± SD based on experimental replicates (n=4) with individual data points showing technical replicates. Technical replicates from each experiment are visualised in the same shade of green/blue, unless indicated differently.

## Results

### Detection of methylglyoxal in the mouse cortex *in vivo*

Mice received an intravenous injection of 25 μg MGO after which brain tissue was collected at time points ranging from 10 minutes to 4 hours after injection. We observed no significant effect of intravenous MGO injection on MGO content in the brain (Fig. 1).

**Figure 1.**
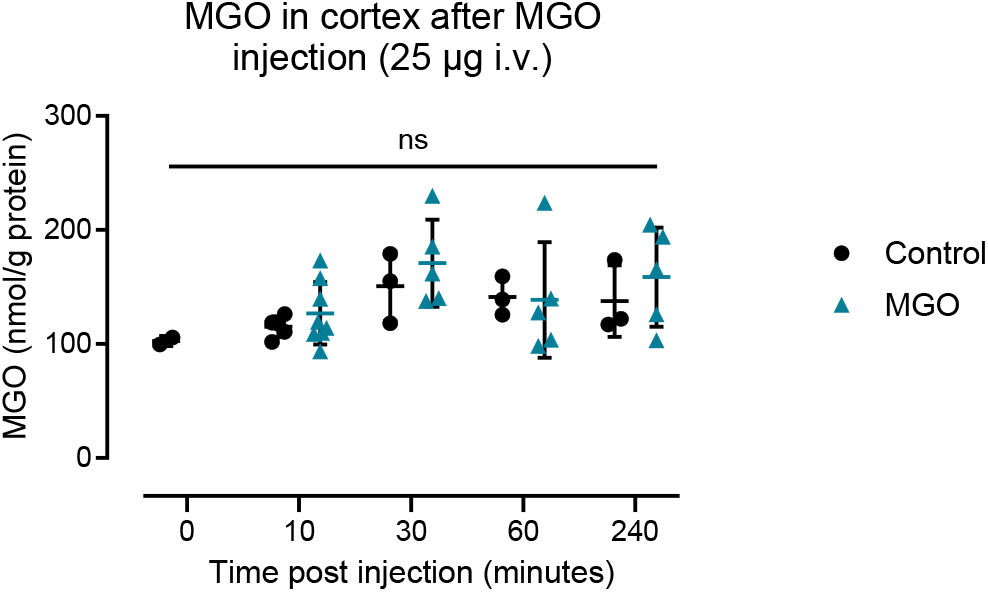
Methylglyoxal (MGO) measured in mouse cortex after 10 minutes, 30 minutes, 1 hour, and 4 hours after intra venous (i.v.) injection of 25 μg MGO (blue triangles) or saline control (black circles). Data in graph represents mean ± SD. Two-way ANOVA: p_time_ = 0.05, p_MGO_=0.26; followed by Tukey’s multiple comparison: not significant (ns).

### Establishing an experimental setup suitable for testing methylglyoxal in vitro

As we could not confirm the passage of MGO into the brain in vivo (Fig. 1), we continued with in vitro BBB models. We assessed the toxicity of MGO and investigated the permeability of highly purified MGO across an in vitro model of the BBB and its possible effect on barrier properties. We exposed the immortalised human brain endothelial cells, hCMEC/D3 cells, to different MGO concentrations in serum free medium ranging from 100 nM to 10 mM for up to 6 hours and investigated cell death with a live/dead staining. We found no significant increase in cell death nor total cell number (dead plus alive) in MGO treated cells compared to the control nor over time (Fig. 2A and 2B).

**Figure 2.**
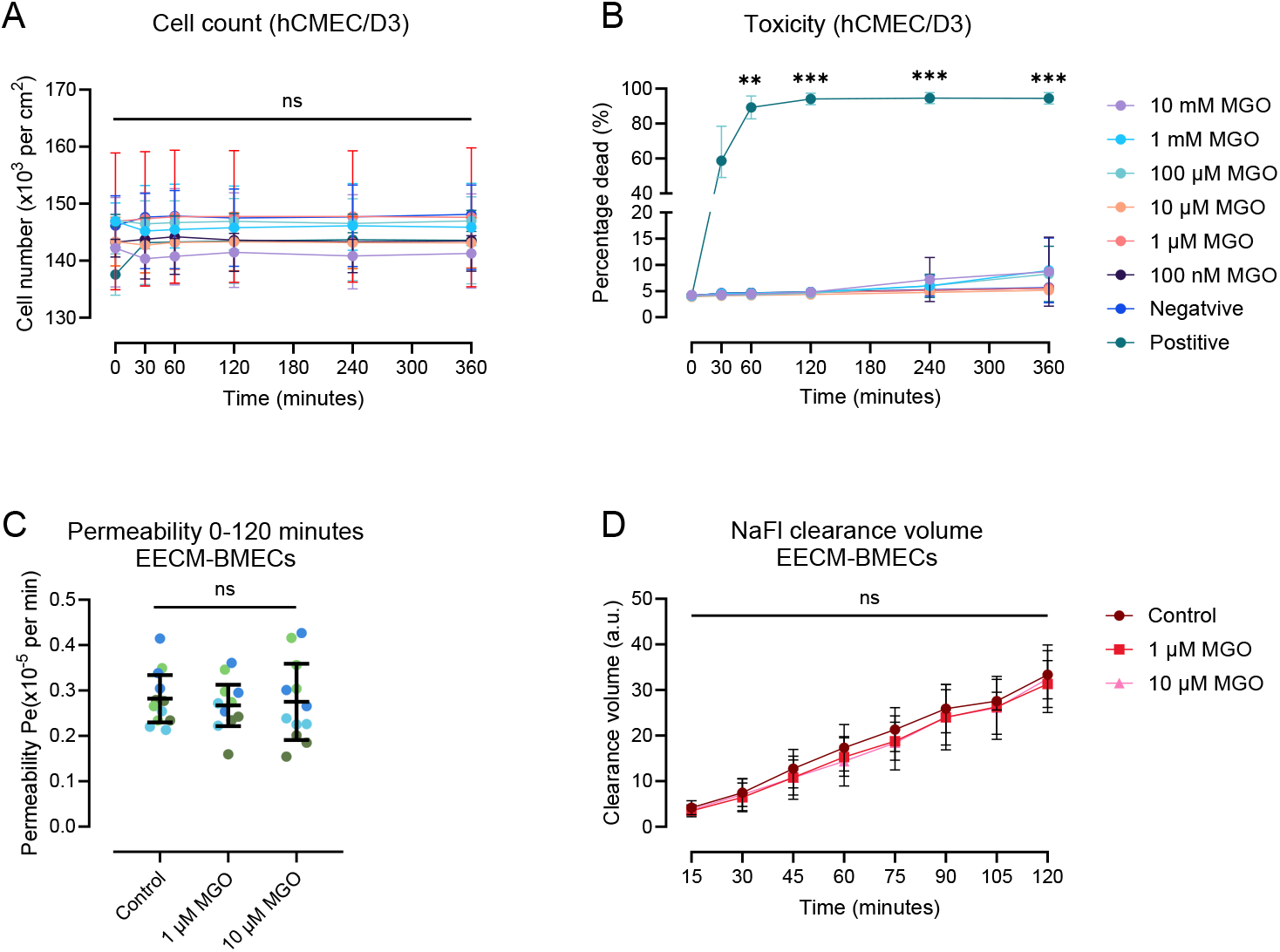
Effect of methylglyoxal (MGO) on cell viability of human cortical microvascular endothelial cells (hCMEC/D3) (A and B) and on cell permeability of EECM-BMECs (C and D). Total number of cells per cm^2^ (A) and percentage of dead cells (B) after exposure to MGO or positive control (10% DMSO) before treatment (0 hours), and after 30 minutes, 1 hour, 2 hours, 4 hours and 6 hours. Cell permeability (Pe) during two hours of MGO exposure or control (C) and the sodium-fluorescein (NaFl) signal in the bottom compartment during the permeability assay over time (D) in EECM-BMECs. The (negative) control in this figure refers to cells exposed to solvent (H_2_O) without additional MGO. Graph data presents mean ± SD, A-B each point represents an average of 3 experiments with 6-8 technical replicates each, C-D n=4 experiments with 2-3 technical replicates each. **p<0.01, ***p<0.001 positive vs negative control (two-way ANOVA, with Dunnett’s multiple comparison).

We found that the MGO concentration in hCMEC/D3 conditioned serum free EBM™-2 medium was 315 ± 90 nmol/L (Fig. S1). Supplementation of 1 μM MGO leads to a 2-fold increase in total MGO concentration, mimicking postprandial MGO spikes observed in humans [20]. To additionally include a supra-physiological MGO concentration, we also included 10 μM MGO which led to a 12-fold increase in MGO (Fig. S1). We therefore used 1 and 10 μM MGO in the experiments from this point onward.

We next studied whether MGO affects BBB properties in hiPSC-derived brain microvascular endothelial cells, as this well-established protocol shows more robust barrier properties compared to the immortalised hCMEC/D3 cell line [17]. EECM-BMECs incubated with 1 or 10 μM MGO for 2 hours showed no increased barrier permeability for NaFl compared to control (Fig. 2C). In EECM-BMECs conditioned hECSR medium, the MGO concentrations were lower (167 ± 7 nmol/L). With addition of 1 and 10 μM MGO, we achieved an approximate 1.6- and 4.3-fold increase in MGO concentration 1 hour after exposure, respectively (Fig. S2). We furthermore studied the NaFl clearance volume for the different treatment groups over time (Fig. 2D). We did not find any difference in NaFl clearance volume over time by MGO exposure (Fig. 2D). These findings showed that 1 or 10 μM of MGO were not toxic to EECM-BMECs and did not impair their barrier properties.

### Methylglyoxal’s passage across the blood-brain barrier in vitro

Next, we investigated whether MGO can cross the EECM-BMEC monolayer. The monolayer was exposed to 1 or 10 μM ^13^C_3_-MGO for 1 hour and the medium was collected from the top compartment (“luminal side”) and bottom compartment (“abluminal side”), and the cell monolayer on the membrane was lysed (figure 3A). Of the total ^13^C_3_-MGO added, we recovered approximately 13% of the ^13^C_3_-MGO in its free form from all the compartments combined after 1 hour exposure, which was the same for all conditions (Fig. 3B). Approximately 1% of the ^13^C_3_-MGO was detected in the bottom compartment of the monolayer for both concentrations, but this amount was significantly lower compared to the control (empty Transwell without cells with 1 μM) (Fig. 3C).

**Figure 3.**
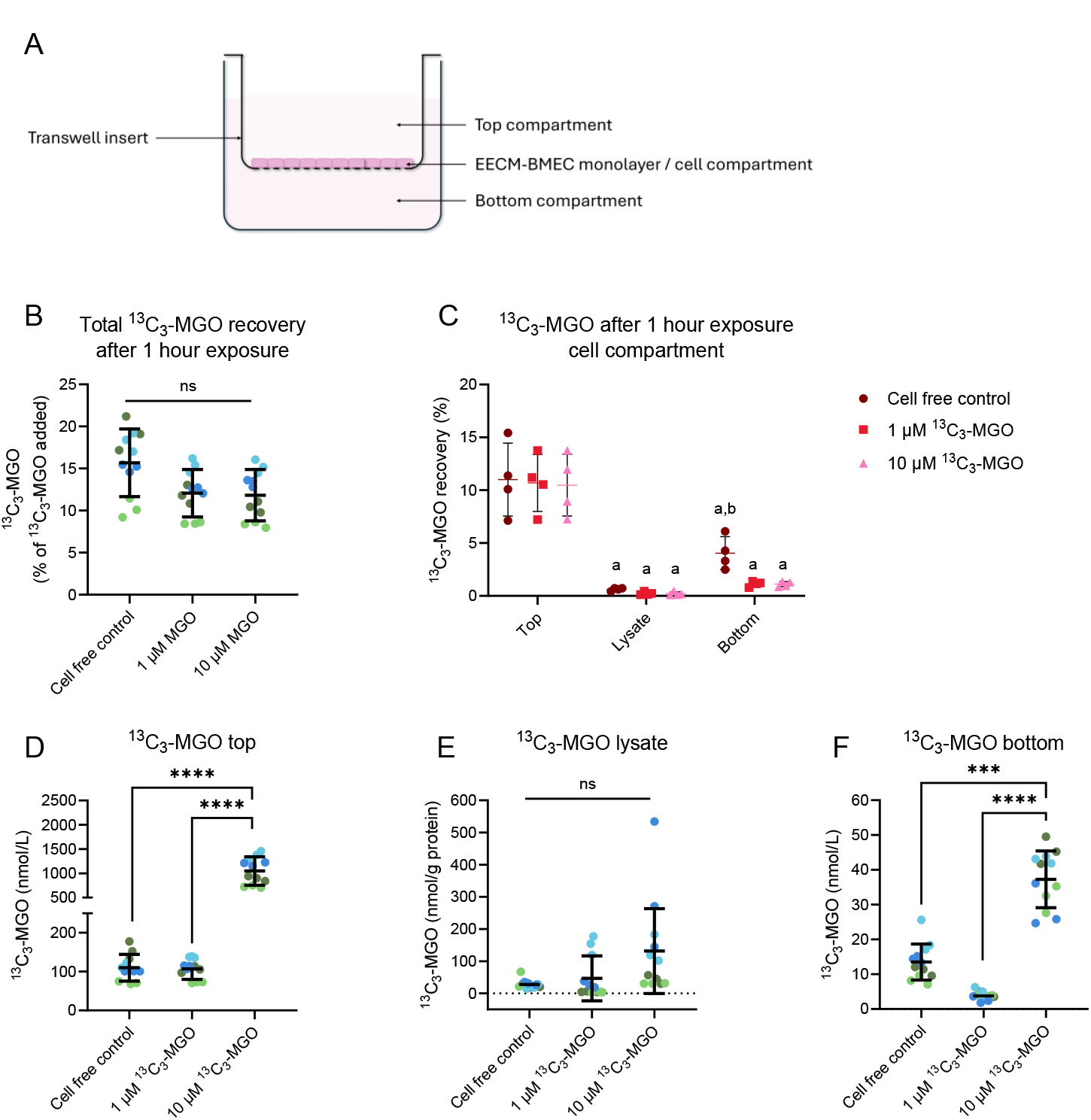
Detection of carbon labelled MGO (^13^C_3_-MGO) in top, cell, and bottom compartments of the *in vitro* model of cells as schematically visualised in panel A. The control represents a cell free Transwell exposed to 1 μM ^13^C_3_-MGO. The percentage of ^13^C_3_-MGO recovered in the entire model (B) and in the different total compartments of the in vitro model after 1 hour incubation (C). The concentration of ^13^C_3_-MGO detected in the top compartment (D), cell monolayer lysate (E), and the bottom compartment (F). Graphs B and D-F present mean ± SD of four experiments (n=4) and each point presents an individual replicate (3 replicates per experiment), one-way ANOVA: *p<0.05, ***p<0.001, ***p<0.0001. Graph C presents mean ± SD of four experiments (n=4), two-way ANOVA: p_compartment_<0.0001, Tukey post-test: ^a^p<0.0001 vs. top, ^b^p<0.05 vs. lysate.

Although the ^13^C_3_-MGO concentration in the top and bottom compartment was approximately 10-fold higher in the 10 μM compared to the 1 μM treated cells and the control (Transwell with 1 µm ^13^C_3_-MGO, but no cells) (Fig. 3D and 3F), there was no significant difference in ^13^C_3_-MGO concentrations in the cell compartment (Fig. 3E). Un-labelled MGO concentrations did not significantly change as a result of MGO exposure in the top and bottom compartment, however, we found that un-labelled MGO measured in the cell lysate was increased in the 10 μM ^13^C_3_-MGO treated cells compared to the 1 μM ^13^C_3_-MGO treated cells and the control (p<0.01) (Fig. S3).

## Discussion

MGO appears to be associated with neurodegenerative disease, however, it remains unclear whether MGO from the circulation can directly cross and/or damage the BBB. To investigate whether MGO can cross the BBB, we measured MGO in the brain cortex of mice following intravenous injection of MGO. Previous experiments using the same dose demonstrated that an increase in MGO in plasma was no longer detectable 10 minutes after injection [16]. We found no significant changes in MGO in the cortex several time points after injection. The absence of a significant increase of MGO in the cortex in MGO injected mice compared to the control may be explained by several factors. First, MGO may not cross the BBB and is therefore not elevated in the brain parenchyma after an MGO bolus injection. Second, MGO might cross the BBB but is rapidly detoxified—e.g., via the glyoxalase system—or bound to proteins, preventing its detection in its free form. Third, if MGO does cross the BBB, its increase might be very localised and undetectable in whole cortical tissue. To be able to more clearly look at the passage of MGO at the brain vasculature, we used an in vitro BBB model.

Remarkably, we did not find any toxic effects of MGO to brain microvascular endothelial cells. This finding is in apparent contrast to previously published in vitro data obtained in BMEC cell lines [21– 24]. In most of these studies, MGO showed a toxic effect in BMEC cells. The discrepancy can be attributed to key differences in experimentally design. Notably, we have used non-contaminated MGO. This could indicate that the contaminants, such as formaldehyde, present in commercially available MGO are responsible for toxic effects reported previously [12]. Furthermore, we have used serum free medium during the MGO exposure to minimise AGE formation and ensure free MGO exposure. This is a critical distinction with previous studies, as MGO is highly reactive in the presence of proteins in serum, significantly reducing availability of free MGO within hours [19]. It must be acknowledged that the absence of serum in the culture medium does not reflect the in vivo situation. However, using serum free medium was an effective way to expose cells to free MGO at a level comparable to the circulation, which was of importance to minimise the possible effects of MGO-derived AGEs. Nevertheless, we conclude that under controlled conditions, free MGO, up to a concentration of 10 mM, is not toxic to brain microvascular endothelial cells in vitro.

To study whether MGO directly damages the BBB, it is of interest to make use of concentrations that are within range of what is observed in the in vivo situation. Previously we have shown that MGO levels in plasma increase approximately 1.5-fold in individuals with a normal glucose metabolism status and approximately 1.7-fold in people with type 2 diabetes after oral glucose load 30 minutes to 1 hour after an oral glucose load [20]. Hence, we have chosen to use 1 and 10 μM additional MGO in our experiments as we showed this led to a 1.6-to-4.3-fold increase, respectively, in medium of the EECM-BMECs one hour after exposure. We found that at these physiological concentrations, MGO did not lead to an increase in barrier permeability.

To study the passage of MGO across the BBB, we made use of labelled MGO to be able to distinguish between MGO added to the luminal side, and MGO naturally present in cells and culture medium. We observed that only 1% of the added labelled MGO was found back in the free form on the abluminal side, which indicates that MGO can cross the endothelium, although minimally. In the cell free control (insert only, no cells), relatively more MGO was measured in the bottom compartment compared to the conditions containing endothelial cells. This suggests that the endothelium reduces the passive flow of MGO from the luminal to the abluminal side. Of note, the overall recovery of labelled MGO was low, which is most likely due to the high reactivity of MGO with residual proteins in the medium or extracellular and membrane proteins produced by the cells. Moreover, we cannot exclude the possibility that part of the MGO is detoxified within the short exposure. Even so, we showed that MGO crosses the BBB, but BMECs prevented or reduced passive flow of MGO, as only a small fraction crossed the brain endothelium in its free form.

While MGO showed no toxic effect on BMECs and did not alter barrier function, a small but significant increase in unlabelled MGO in the cell lysate of cells treated with a high concentration of MGO suggests a saturation and/or impairment of the detoxification system in response to MGO addition. The consequence of repeated exposure to very high MGO concentrations warrants further investigated.

The use of well-characterised in vitro BBB model allowed opportunities to study the passage of MGO across the endothelium. However, some limitations of this model include the absence of other cell types, such as pericytes and astrocytes, and the lack of fluid flow, which could strongly affect the dynamics of MGO with the brain endothelium. Moreover, astrocytes, an important player in the BBB in vivo, are known to be strong detoxifiers of MGO [25] and would in theory be able to detoxify the small fraction of MGO that crosses. We speculate that, while MGO crosses the brain microvascular endothelium in very small amounts, the effect on brain function is likely minimal.

To get a better understanding of what the consequences of the small transit of MGO might be after long-term exposure, it would be of interest to investigate the glycation and detoxification of MGO. Furthermore, a limitation of our study is that we only made use of healthy in vivo and in vitro conditions, while it is known that during disease MGO levels in plasma are increased and BBB integrity is reduced [5, 7, 26, 27]. It has been suggested that an increase in MGO in the brain predominantly occurs during age-related pathology [28]. Therefore, these experiments should be repeated in models of disease.

We aimed to determine whether MGO can damage the endothelial cells which could cause BBB breakdown. In summary, we found that MGO does not damage brain microvascular endothelial cells or impair BBB barrier function. We furthermore showed that free MGO can cross the BBB, but the amount is very low. In conclusion, extracellular MGO is less toxic to the BBB as previously shown, and its limited passage across the BBB implies that plasma MGO is unlikely to have direct effects on neuronal functioning under normal physiological conditions.

## Author contributions

EB – Conceptualization, Investigation, Formal analysis, Writing – original draft. SG – Investigation, Methodology, Writing – reviewing and editing. XZ – Resources, Writing – reviewing and editing IF – Investigation. MvdW – Investigation. JS – Methodology, Investigation, Resources. KW – Resources, writing – Reviewing and editing. RvO – Supervision, Funding acquisition, Writing –Reviewing and editing. BE – Methodology, Resources, Supervision, Writing – Reviewing and editing SF – Conceptualization, Funding acquisition, Supervision, Writing – review and editing. CS –Conceptualization, Funding acquisition, Supervision, Writing – review and editing. All authors have seen the manuscript and have approved to submit the manuscript.

## Funding

This study was funded by EFSD/Boehringer Ingelheim European Research Programme in Microvascular Complications of Diabetes 2018, the Harry Struijker-Boudier Award For Talented Academics (HS-BAFTA) Talented PhD Candidates grant from the Cardiovascular Research Institute Maastricht (CARIM), and additional funding from CARIM.

## Notes

### Competing Interest Statement

The authors have declared no competing interest.

