## Supplementary data for "Extracellular methylglyoxal; the passage across brain endothelial cells and the effect on barrier function"

* Shared last-authorship

**Affiliations**

[1] Department of Internal Medicine, Maastricht University, Maastricht, the Netherlands.

[2] Cardiovascular Research Institute Maastricht (CARIM), Maastricht University, Maastricht, the Netherlands.

[3] Theodor Kocher Institute, University of Bern, Bern, Switzerland

[4] Department of Neurology, Maastricht University Medical Centre, Maastricht, the Netherlands.

[5] MHeNs, Mental Health and Neurosciences Research Insitute, Maastricht University, Maastricht, the Netherlands.

[6] Department of Pharmacology and Toxicology, Maastricht University, Maastricht, the Netherlands.

**Corresponding author**

Prof Dr Casper Schalkwijk

Maastricht University, Faculty of Health Medicine and Life Sciences

Department of Internal Medicine

Universiteitssingel 50, 6229 ER Maastricht, The Netherlands

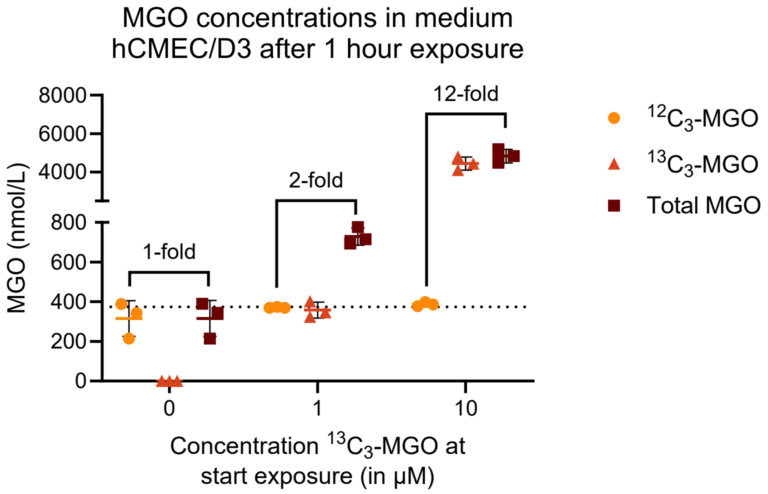


**Supplemental figure 1.** Methylglyoxal (MGO) concentrations in medium after $^{13}C_3$-MGO exposure of hCMEC/D3 cells. Circles represent the non-labelled $^{12}C_3$-MGO present in medium exposed to cells, triangles represent the recovered free labelled $^{13}C_3$-labelled MGO after 1 hour of exposure, and squares represent the total (labelled + non-labelled) MGO present in the medium 1 hour after exposure. Bars show mean ± SD, the dotted line presents the median concentration of non-labelled MGO present in medium, each data point is a technical replicate within one experiment.

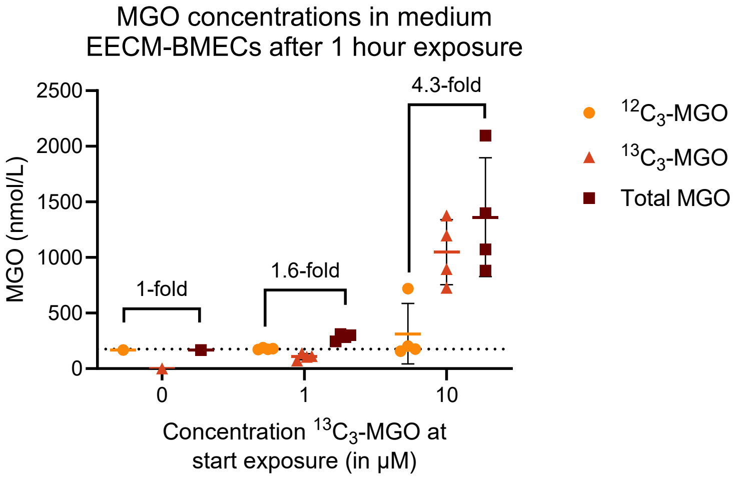


**Supplemental figure 2.** Methylglyoxal (MGO) concentrations in medium after $^{13}C_3$-MGO exposure of EECM-BMECs (B). Circles represent the non-labelled $^{12}C_3$-MGO present in medium exposed to cells, triangles represent the recovered free labelled $^{13}C_3$-labelled MGO after 1 hour of exposure, and squares represent the total (labelled + non-labelled) MGO present in the medium 1 hour after exposure. Bars show mean ± SD, the dotted line presents the median concentration of non-labelled MGO present in medium. Each data point represents an experimental replicate using the mean of three technical replicates.

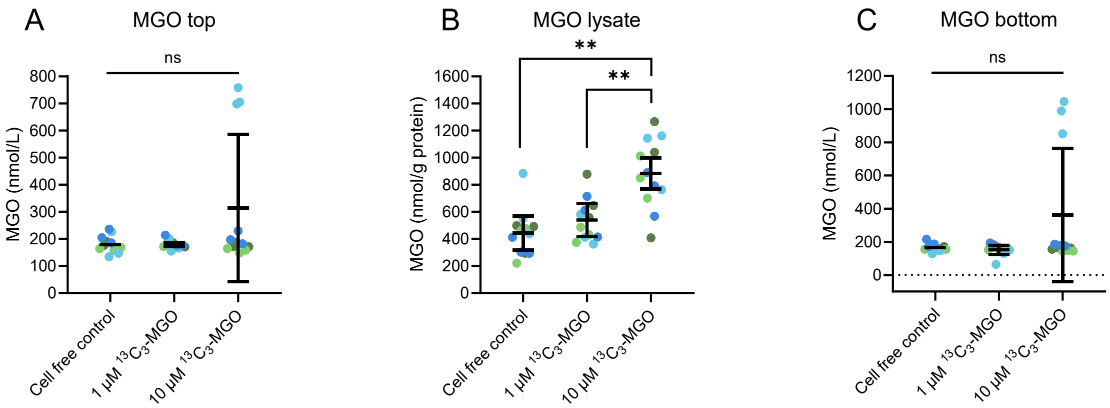


**Supplemental figure 3.** Non-labelled methylglyoxal ($^{12}C_3$-MGO) in the top (A), cell monolayer lysate (B), and the bottom compartment (C). The cell free control represents a Transwell insert without cells exposed to 1 µM $^{13}C_3$-MGO. Graphs present mean ± SD of four experiments (n=4) and each point presents an individual replicate (3 replicates per experiment), one-way ANOVA: **$p<0.01$ or non-significant (ns).
